# Signposts in the fog: objects facilitate scene representations in left scene-selective cortex

**DOI:** 10.1101/206003

**Authors:** Talia Brandman, Marius V. Peelen

## Abstract

We internally represent the structure of our surroundings even when there is little layout information available in the visual image, such as when walking through fog or darkness. One way in which we disambiguate such scenes is through object cues; for example, seeing a boat supports the inference that the foggy scene is a lake. Recent studies have investigated the neural mechanisms by which object and scene processing interact to support object perception. The current study examines the reverse interaction, by which objects facilitate the neural representation of scene layout. Photographs of indoor (closed) and outdoor (open) real-world scenes were blurred such that they were difficult to categorize on their own, but easily disambiguated by the inclusion of an object. fMRI decoding was used to measure scene representations in scene-selective parahippocampal place area (PPA) and occipital place area (OPA). Classifiers were trained to distinguish response patterns to fully visible indoor and outdoor scenes, presented in an independent experiment. Testing these classifiers on blurred scenes revealed a strong improvement in classification in left PPA and OPA when objects were present, despite the reduced low-level visual feature overlap with the training set in this condition. These findings were specific to left PPA/OPA, with no evidence for object-driven facilitation in right PPA/OPA, object-selective areas, and early visual cortex. These findings demonstrate separate roles for left and right scene-selective cortex in scene representation, whereby left PPA/OPA represents inferred scene layout, influenced by contextual object cues, and right PPA/OPA represents a scene’s visual features.

## Introduction

We recognize scenes at a glance, even though they contain rich and complex visual information (Potter, 1975). The ability to rapidly categorize scenes (e.g., as indoor or outdoor) has been shown to depend on scene-selective regions in visual cortex - regions defined by stronger neural responses to scenes and buildings than to isolated objects (Aguirre, Zarahn, & D’Esposito, 1998; R. Epstein & Kanwisher, 1998). For example, when activity in the scene-selective occipital place area (OPA) is disrupted by TMS, participants are less accurate in scene categorization, while object categorization remains unaffected (Dilks, Julian, Paunov, & Kanwisher, 2013; Ganaden, Mullin, & Steeves, 2013). This supports the distinction between object- and scene-selective pathways (Harel, Kravitz, & Baker, 2013; Mullin & Steeves, 2011; Park, Brady, Greene, & Oliva, 2011). In everyday life, however, scenes and objects are perceived together and their processing heavily interacts, as observed in behavioral studies (Bar & Ullman, 1996; Biederman, Mezzanotte, & Rabinowitz, 1982; Davenport & Potter, 2004; Munneke, Brentari, & Peelen, 2013; Oliva & Torralba, 2007). How are these interactions implemented in visual cortex?

In a recent study, we tested how scene processing in scene-selective cortex biases object processing in object-selective cortex (Brandman & Peelen, 2017). In that study, objects were hard to recognize on their own but easy to recognize when presented within their original scene context. This scene-based disambiguation of object processing was reflected in more distinct multivariate activity patterns in object-selective areas, with the strength of this effect being predicted by activity in scene-selective areas (Brandman & Peelen, 2017). In the current fMRI study, we investigated the reverse interaction, testing how object processing disambiguates scene perception. Similar to how scene context disambiguates object perception, the presence of objects allows us to interpret an otherwise ambiguous scene (e.g. Figure 1a), as often happens to us in darkness or fog, in which layout information is veiled. To our knowledge, this role of objects in scene perception has not been explored. Thus, to test for influences of object processing on scene representation, we examined the neural representation of scene category (indoor, outdoor) in blurred scenes, which were difficult to categorize on their own but easily disambiguated by inclusion of an intact object.

**Figure 1:**
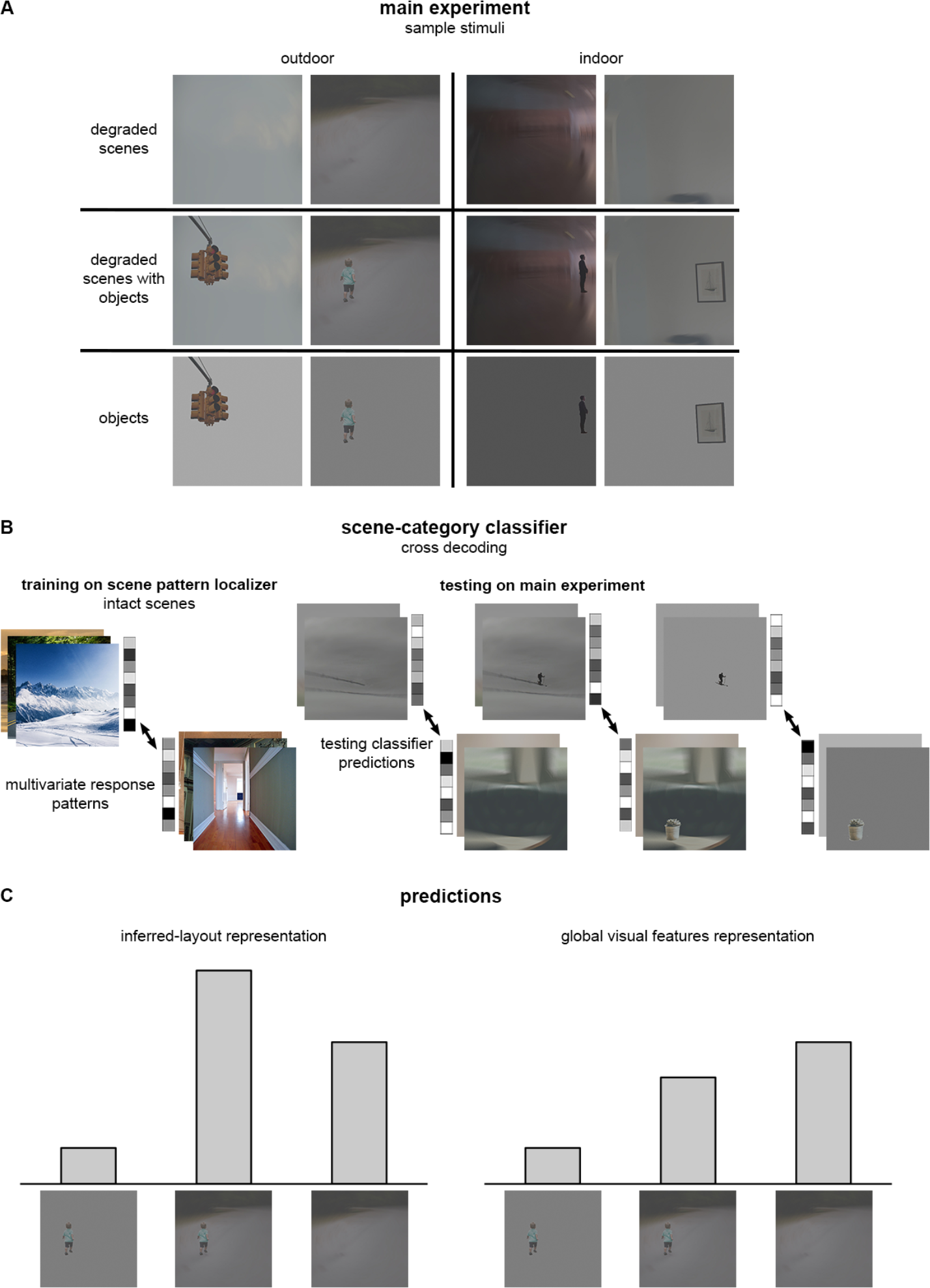
Experimental design and predictions. (A) Sample stimuli of the main experiment conditions: indoor (closed) and outdoor (open) degraded (blurred) scenes, degraded scenes with objects and objects on gray background of mean luminance of the degraded scene; (B) Cross-decoding analysis, whereby a classifier was trained on indoor/outdoor scene discrimination from the fMRI response patterns, and tested on discrimination of indoor/outdoor degraded scenes, degraded scenes with objects, and isolated objects; (C) Prediction of cross-decoding results (decoding accuracy) in areas representing inferred scene layout (left panel) and in areas representing scenes’ global visual features (right panel).

In addition to revealing interactions between object- and scene-selective pathways, the current study addresses a recent debate about the representational content of scene-selective areas. With their discovery, the critical factor in scene-selective responses was found to be the presence of spatial layout information. It was therefore suggested that scene-selective areas represent place information by encoding the geometry of the local environment (R. Epstein & Kanwisher, 1998), as part of a network involved in spatial navigation (R. A. Epstein, 2008). One source of information about the spatial layout of scenes (e.g., open vs closed) is provided by global visual features, with second-order image statistics being informative for scene category or scene “gist” (Torralba & Oliva, 2003). This raises the possibility that spatial layout information in scene-selective areas (Kravitz, Peng, & Baker, 2011; Park et al., 2011; Walther, Caddigan, Fei-Fei, & Beck, 2009) reflects the feedforward processing of such visual features. This view is in line with recent studies showing that scene selectivity itself can be (partly) explained by sensitivity to relatively low-level visual features such as cardinal orientations and rectilinearity (Nasr, Echavarria, & Tootell, 2014; Nasr & Tootell, 2012; Rajimehr, Devaney, Bilenko, Young, & Tootell, 2011; Zeidman, Mullally, Schwarzkopf, & Maguire, 2012). Alternatively, scene representations in scene-selective areas may be more abstract, representing the layout of a scene as inferred based on all cues available to the individual (R. A. Epstein, 2008; Peelen & Downing, 2017; Wolbers, Klatzky, Loomis, Wutte, & Giudice, 2011).

Here we distinguish between these two accounts by measuring the level of scene disambiguation gained by object cues (Figure 1c). These objects add contextual cues without adding visual features associated with spatial layout. We therefore predicted that regions representing global visual scene features should carry most information about scenes without objects, because objects would only act as clutter. By contrast, regions representing inferred spatial layout should carry most information about scenes in which the layout information is disambiguated by the objects.

To measure the amount of scene-category information gained by the inclusion of object cues, photographs of real-world indoor (closed) and outdoor (open) scenes were blurred such that they were difficult to categorize on their own but easily categorized with an intact object overlaid on the scene in its original position. In this way, scene category (or scene layout), processed in scene-selective areas (Kravitz et al., 2011; Park et al., 2011; Walther et al., 2009), was disambiguated by object cues. The objects were also presented alone, to assess the baseline level of scene-category information carried by the objects themselves (Harel et al., 2013). We then measured scene-category information using multivariate pattern analysis (MVPA; Haxby et al., 2001), with classifiers trained on the response patterns evoked by intact scenes in a separate experiment, and tested on responses evoked by degraded scenes, degraded scenes with objects, and objects alone (Figure 1b). Within each tested brain region, the effect of objects on contextual disambiguation of scenes was measured by the difference in decoding accuracy for degraded scenes with and without objects. Thereby, we were able distinguish between areas representing inferred scene layout and areas representing visual scene features.

## Methods

For degraded scenes, degraded scenes with objects, and for objects alone, we measured the multivariate representations of scene category in the fMRI signal while participants performed a 1-back task (see Procedure). In a separate *pattern localizer* used for classifier training, participants viewed intact indoor and outdoor scenes, fully visible and in high resolution, presented without objects. In addition, a region-of-interest (ROI) localizer served to localize scene- and object-selective ROIs in visual cortex. All procedures were approved by the ethics committee of the University of Trento.

### Participants

Nineteen healthy participants (8 female, mean 24 years ± 2.98 SD) were included. All participants had normal or corrected to normal vision and gave informed consent. Sample size was chosen to match that of previous work examining contextual effects of object and scene integration with similar fMRI decoding methods (Brandman & Peelen, 2017). Six additional participants were excluded from data analysis due to excessive head motion during scanning (n=4), widespread anatomical artefact (n=1), or failure to follow task instructions (n=1).

### Stimuli

The stimulus set consisted of degraded scenes that were perceived as ambiguous on their own but that were easily categorized when presented with an object. Both indoor and outdoor scenes included a mixture of animate and inanimate objects of various categories. We ensured that the scenes did not contain other objects contextually associated with the foreground objects. The main experiment included photographs of 30 indoor and 30 outdoor scenes. Sixty photographs of scenes from Unsplash (unsplash.com) and Pixabay (pixabay.com) were cropped and cleaned to include one dominant foreground object. The scene excluding the object was degraded (blurred) and contrast was reduced for the entire image. Scenes were saved with and without the object (object was edited out). The object was also saved in isolation, on a uniform gray background of mean luminance of the original background. The final images (180 in total) included the degraded scenes, degraded scenes with objects and objects alone (see samples in Figure 1a). To avoid familiarity effects passing from scenes with objects to scenes alone, the stimulus set was split in two, such that different scenes were presented for degraded scenes with objects and for degraded scenes alone within a given subject (For example, for two stimuli Beach1 and Beach2, a given participant would either view Beach1 alone and Beach2 with a boat, or vice versa). The objects alone matched the degraded scenes alone (participants who viewed Beach1 alone would also view its boat separately, but not embedded). The two sets were counterbalanced across subjects. The pattern localizer used included the 60 scenes from the main experiment (with different cropping), and an additional set of 60 new scenes that were matched for category and sub category of the main-experiment set, in high resolution. Visual angle: 5.99x5.99 degrees (400×400 pixels).

### Stimulus Optimization and Selection

The stimulus set was optimized and validated in an online behavioral pilot experiment in Amazon Mechanical Turk, in which participants rated the degraded scenes’ category on a scale of indoor-outdoor, to compare the level of scene ambiguity with and without objects. Participants were asked to rate each degraded scene, presented either with its original embedded object or alone, on a scale from 1 - indoor to 8 - outdoor. Participants’ ratings were normalized to a mean of 0 and standard deviation of 1. The final stimulus set, presented in the MRI, consisted of scenes that were perceived as more ambiguous on their own than with objects (N=29; indoor: *t* = 10.83, *p* < 0.001; outdoor: *t* = 15.69, *p* < 0.001), with a mean difference of 1.02 between normalized scores of scenes with and without objects. Forty-five additional scenes, showing smaller differences in ratings (with versus without an object), were tested in the piloting stages but were excluded from the final stimulus set.

### Procedure

On each trial, participants viewed a single briefly presented (80 ms) stimulus. Throughout all runs participants performed a 1-back task in which they pressed a button each time an image appeared twice in a row. The main experiment consisted of 5 scanner runs of 352 s duration, each composed of 4 fixation blocks and 3 blocks for each of the 6 conditions: indoor/outdoor x scene/scene-with-object/object (total 18). The main experiment was followed by a pattern localizer, which consisted of 3 scanner runs of 336 s duration, each composed of 5 fixation blocks and 4 blocks per condition: indoor/outdoor x old/new (total 16). Each block consisted of 16 trials, in which a stimulus was presented for 50 ms followed by a 950 ms fixation. This resulted in 240 trials (120 2-sec volumes) per condition in the main experiment, and 192 trials (96 2-sec volumes) per condition in the pattern localizer. A category-selectivity localizer ended the scanning session, designed to identify scene- and object-selective areas. The localizer included 80 grayscale images per category of objects, scenes, bodies and scrambled objects (i.e. a random mixture of pixels of each of the object images). It consisted of 2 scanner runs of 336 s duration, each composed of 5 fixation blocks and 4 blocks per condition: scene/object/body/scrambled-object (total 16). Each block consisted of 20 trials, in which a stimulus was presented for 350 ms followed by a 450 ms fixation.

### Data Acquisition and Preprocessing

Whole-brain scanning was performed with a 4T Bruker MedSpec MRI scanner using an 8-channel head-coil. T2*-weighted EPIs were collected (TR = 2.0 s, TE = 33 ms, 73° flip angle, 3 × 3 × 3 mm voxel size, 1-mm gap, 30 slices, 192-mm FOV). A high-resolution T1-weighted image (magnetization prepared rapid gradient echo; 1 × 1 × 1 mm voxel size) was obtained as an anatomical reference. The data were analyzed using MATLAB (MathWorks) with statistical parametric mapping (SPM). Each run was preceded by 12 s fixation discarded from the analysis. Preprocessing included slice-timing correction, realignment and spatial smoothing with a 6 mm full-width at half-maximum (FWHM) Gaussian kernel. A GLM HRF model was estimated for each participant for the univariate analyses.

### Category-selective ROIs

Functional ROIs were defined using a two-step group-constrained subject-specific method (e.g. Julian, Fedorenko, Webster, & Kanwisher, 2012). The first selection step was based on results of a group analysis in MNI space. Scene-selective areas were defined by contrasting activity evoked by scenes against objects, against scrambled objects, and against baseline activity. The parahippocampal place area (PPA) and occipital place area (OPA) ROIs were generated by identifying temporal and occipital voxels in the ventral visual stream where all 3 contrasts garnered uncorrected p values less than 0.01 at group-level (random effects). The retrosplenial complex (RSC) was too small to perform MVPA (<15 voxels), and was therefore excluded from the analysis. Similarly, object-selective areas were defined by contrasting activity evoked by objects against scenes, against scrambled objects, and against baseline activity. The posterior fusiform sulcus (pFs) and lateral occipital (LO) ROIs were generated by identifying temporal and occipital voxels in the ventral visual stream where all 3 contrasts garnered uncorrected p values less than 0.01 at group-level (random effects). The second ROI-selection step was performed for each participant separately, where group-selected ROIs were used as inclusion masks for individual ROI selection. Only the most significant 50 voxels of each ROI, as measured by individually-estimated T-values, were selected for multivariate analysis.

### Early visual areas

Early visual ROIs were defined separately for each participant, in each hemisphere, by selecting the most significant 50 voxels as measured by an individually-estimated T-contrast of scrambled objects against baseline activity, within Brodmann area 17, and excluding voxels showing higher responses to scenes than scrambled objects.

### Multivariate Analysis

The data within each voxel were detrended and normalized (mean and STD) across the time-course of each run, and shifted two volumes (4 s) to account for the hemodynamic lag. The data were then averaged across blocks within each run, resulting in one block of 8 volumes per condition per run. Multivariate analysis was performed using the CoSMoMVPA toolbox (Oosterhof, Connolly, & Haxby, 2016). An SVM classifier discriminated between response patterns to indoor vs. outdoor scenes. The decoding approach is illustrated in Figure 1b. First, decoding of intact scene category was measured within the pattern localizer, by training on old-scene trials (i.e. scenes included in the main-experiment set), and testing on new-scene trials. Next, cross-decoding was achieved by training on all conditions of the pattern localizer, and testing on each of the main-experiment conditions (scene, scene-with-object, object). For each participant, cross decoding was performed across the voxels of each ROI, resulting in an overall accuracy score for the ROI for each of the 3 conditions.

### Controlling for multiple comparisons

All significant t-tests and correlations reported remained significant when correcting for multiple comparisons within each section of the Results, using false discovery rate (FDR) at a significance level of .05.

## Results

### Decoding intact scene category in scene-selective cortex

We first assessed the representation of intact scene category (indoor vs outdoor scenes) in scene-selective areas for the pattern localizer data. Results showed that scene category was strongly represented in four scene-selective areas: the right PPA (decoding accuracy *M* = 72.37%;), left PPA (*M* = 72.48%;), right OPA (*M* = 64.80%;) and left OPA (*M* = 68.86%;), and was decoded significantly above chance (*t*(18) > 4.69, *p* < 0.001, for all regions). These results are in line with previous findings of scene-category decoding in visual cortex (Walther et al., 2009). We found no significant effects of region (PPA, OPA; *F*(1,18) = 4.12, *p* = 0.057) hemisphere (right, left; *F*(1,18) = 0.88, *p* = 0.360) or their interaction (*F*(1,18) = 1.22, *p* = 0.284) in intact scene decoding.

### Decoding degraded scene category in scene-selective cortex

Next, we examined the representation of scene-category in scene-selective areas, the left and right PPA and OPA, for each of the three main experiment conditions (Figure 2). Classifiers were trained on data from the pattern localizer and tested on the conditions in the main experiment using a cross-decoding approach (Figure 1b). Importantly, the effect of our context manipulation varied across hemispheres, as revealed by a two-way interaction (*F*(2,36) = 10.35, *p* < 0.001) between hemisphere (right, left) and context (scene, scene with object, object). There was no interaction between region (PPA, OPA) and context (*F*(2,36) = 1.65, *p* = 0.205), and no three-way interaction (*F*(2,36) = 0.30, *p* = 0.741). Considering this pattern of results, we examined the average decoding across PPA and OPA, separately for each hemisphere.

**Figure 2:**
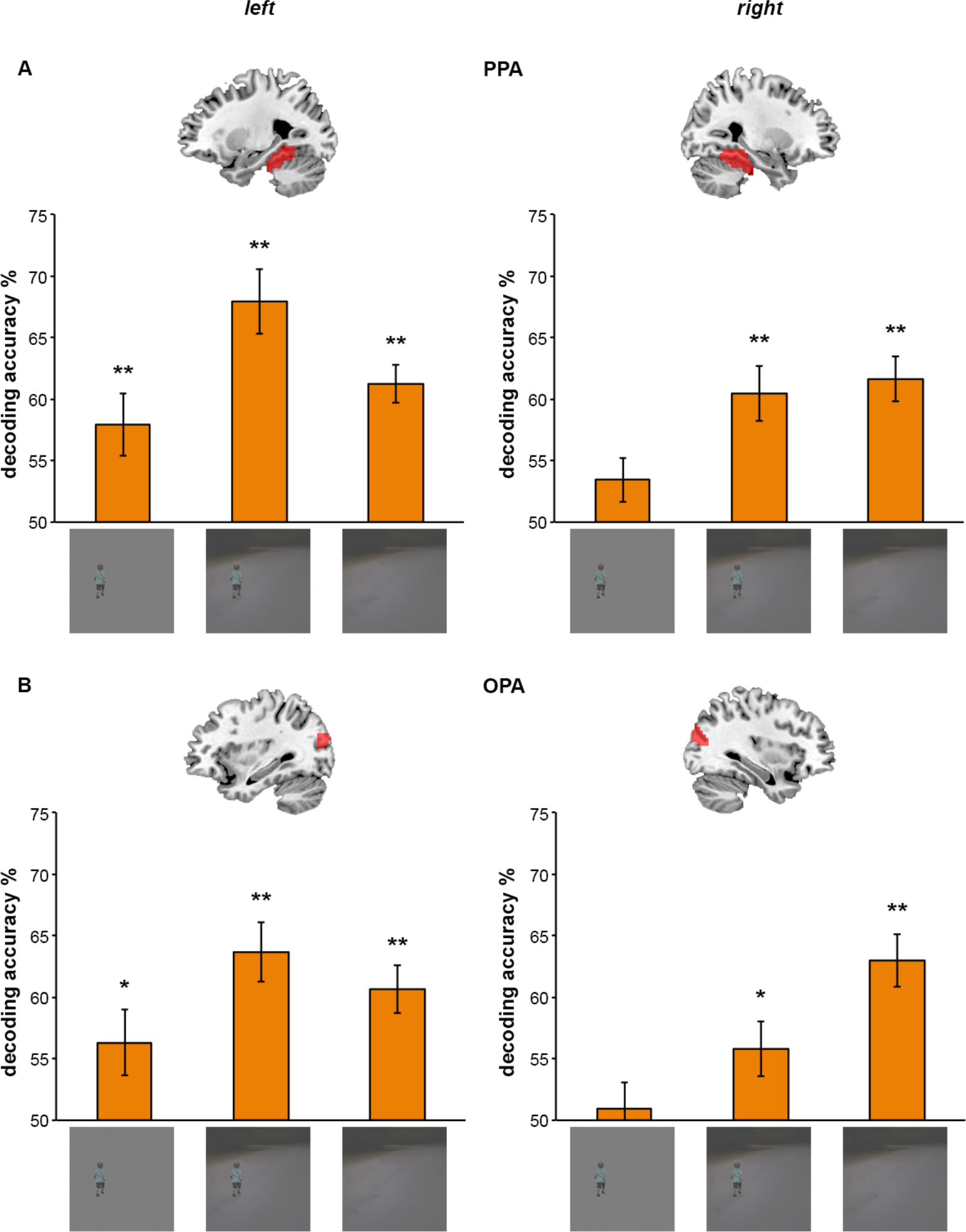
Cross-decoding scene category in scene-selective areas. Scene-selective areas (red regions in brain maps) were defined by stronger responses to scenes than to objects and scrambled objects in a separate localizer run: (A) Parahippocampal place area (PPA); (B) Occipital place area (OPA). In the left hemisphere, decoding of scene layout (indoor – closed, outdoor – open) from degraded scenes was better for degraded scenes with object than without. In the right hemisphere, objects did not inform scene-category decoding. Thus, objects facilitated the classification of degraded scenes in left scene-selective areas, but not right scene-selective areas. Data are represented as mean distance from chance (50% decoding accuracy) ±SEM. \**p*<0.05. \*\**p*<0.01.

The category of degraded scenes could be reliably decoded in both hemispheres of scene-selective areas (against chance; *t*(18) > 7.98, *p* < 0.001, for both hemispheres). However, the role of objects in the representation of scene category varied across hemispheres, corresponding to the two hypothesized response profiles (Figure 1c). Particularly, in left scene-selective areas, objects significantly facilitated the decoding of degraded scenes compared to when the scenes were presented without objects (paired *t*(18) = 2.25, *p* = 0.037). This was not observed in right scene-selective areas, which showed a trend in the opposite direction (paired *t*(18) = 1.80, *p* = 0.089).

### Univariate differences in scene-selective cortex

In addition, to test whether hemispheric differences found in scene-selective areas were related to differences in overall activation in these regions, we examined their indoor and outdoor univariate BOLD responses. Data were processed similarly as for multivariate analysis, excluding the normalization step. There was a main effect of scene category (indoor, outdoor; *F*(1,18) = 19.46, *p* < 0.001) with higher activity for indoor than outdoor scenes, replicating previous reports (Henderson, Larson, & Zhu, 2007). Importantly, scene category did not interact (*F* < 2.52, *p* > 0.094, for all tests) with context (scene, scene with object, object), region (PPA, OPA), or hemisphere (right, left). Thus, the differences found in multivariate representations of scene category across experimental conditions and hemispheres cannot be explained by regional response-magnitude differences.

### Decoding scene category from objects in scene-selective cortex

Interestingly, results showed that objects presented in their original position on a gray background with the mean luminance of the original scene were sufficient predictors of scene category in left scene-selective areas (against chance; *t*(18) = 3.07, *p* = 0.007), but not right scene-selective areas (against chance; *t*(18) = 1.60, *p* = 0.127). We hypothesized that this may reflect a similar (though reduced) perceptual inference as observed in the degraded scene with object condition, with the mean luminance background acting as a more extremely degraded scene. We therefore asked whether object-driven facilitation of blurred scenes was associated, across participants, with object-based decoding of mean-luminance backgrounds. Results revealed that the level of object-driven facilitation, as measured by the difference in decoding accuracies between scenes with and without objects, was indeed highly correlated with decoding of objects on mean luminance backgrounds in left scene-selective areas (*r*(17) = 0.73, *p* < 0.001), but not right scene-selective areas (*r*(17) = 0.39,*p* = 0.100) (Figure 3). These results provide further evidence for the hemispheric specificity of object-based facilitation and suggest a common origin for the effects observed in the scene-with-object and background-with-object conditions.

**Figure 3:**
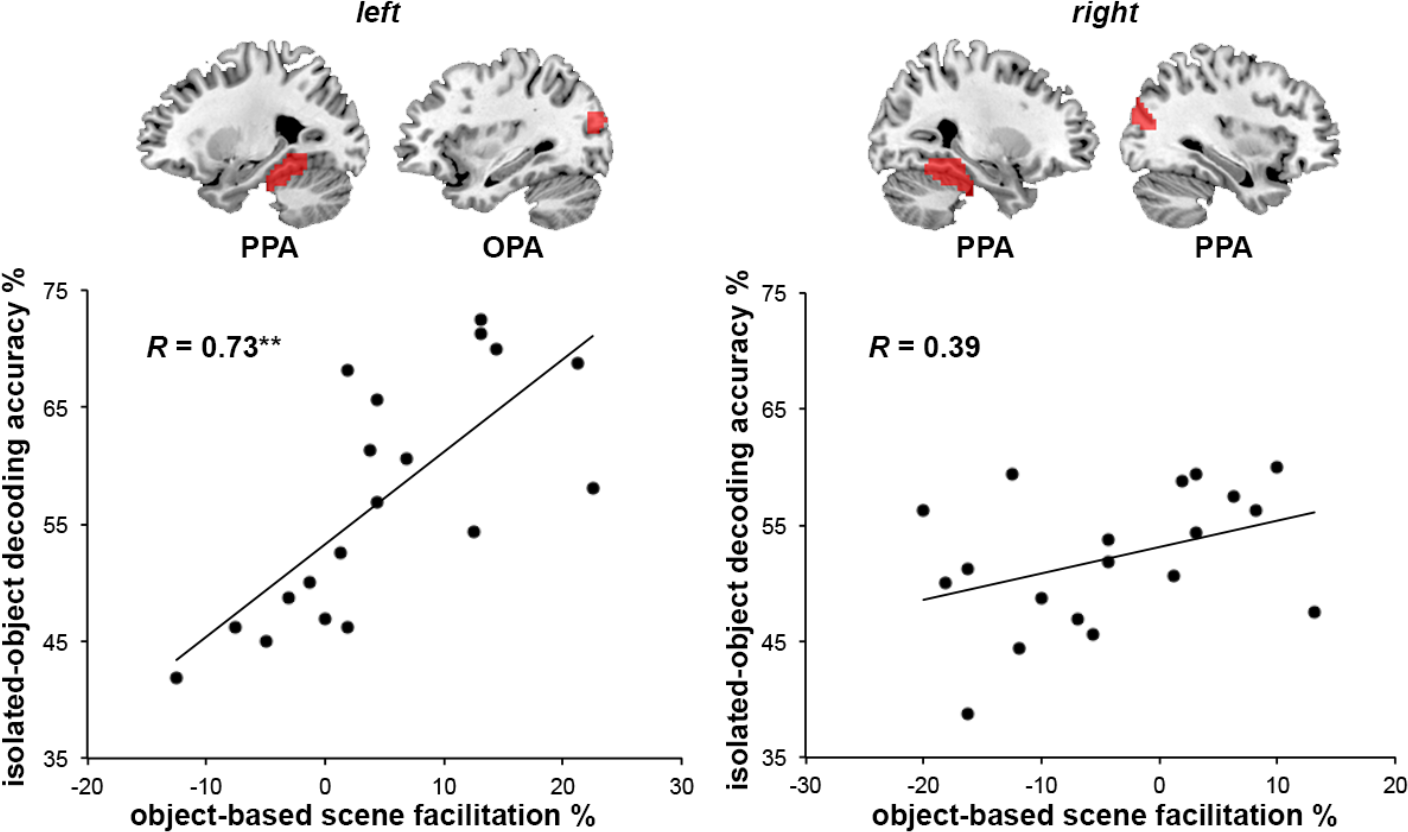
Correlation of object-driven facilitation of scene decoding with object-based scene decoding. The difference in decoding accuracies for degraded scenes with and without objects was significantly correlated with the decoding accuracy for isolated objects in left, but not right, PPA/OPA. \*\**p*<0.01

### Decoding degraded scene category in early visual cortex

To test whether the hypothesized profile of visual scene representation (Figure 1c) correctly predicted low-level visual processing, we examined the decoding of scene category also in early visual cortex (Figure 4), using the same cross-decoding approach as for scene-selective areas (Figure 1b). Indeed, scene category was reliably decoded in early visual cortex (EVC) for scenes alone (against chance; *t*(18) = 3.89, *p* = 0.001) but not for scenes with objects (against chance; *t*(18) = 1.38, *p* = 0.184) or objects alone (against chance; *t*(18) = 0.033, *p* = 0.974). Decoding significantly varied between these three context conditions (*F*(2,36) = 13.62,*p* < 0.001), with no effect of hemisphere (*F*(1,18) = 0.11,*p* = 0.741), and no interaction (*F*(2,36) = 0.61, *p* = 0.551).

**Figure 4:**
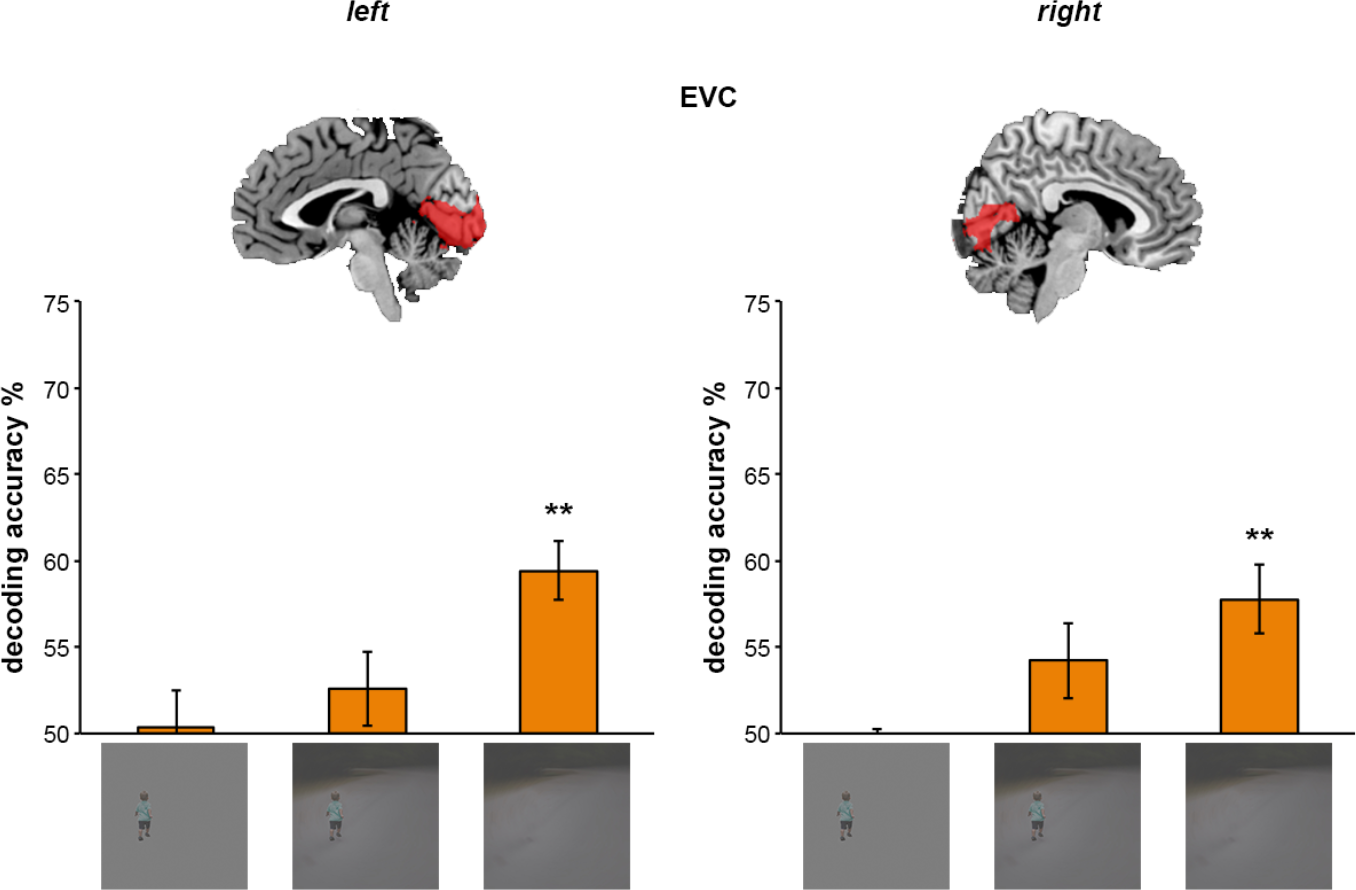
Cross-decoding scene category in early visual areas. Early visual areas (red regions in brain maps) were defined by stronger responses to scrambled objects than baseline activity within Brodmann area 17. Decoding of scene layout (indoor – closed, outdoor – open) was most accurate for degraded scenes alone, and objects did not inform scene decoding. Thus, objects did not facilitate the classification of degraded scenes in early visual areas. Data are represented as mean distance from chance (50% decoding accuracy) ±SEM. \*\**p*<0.01.

### Decoding degraded scene category in object-selective cortex

Finally, we examined representations of scene category in object-selective areas (Figure 5), using the same cross-decoding approach as for scene-selective areas (Figure 1b). Results showed no differences in context (scene, scene with object, object; *F*(2,36) = 2.40, *p* = 0.105), region (LO, pFs; *F*(1,18) = 3.27, *p* = 0.087) or hemisphere (left, right; *F*(1,18) = 0.53, *p* = 0.477), nor any interactions between them (*F* < 1.36, *p* > 0.259). Thus, in contrast to scene-selective areas in the left hemisphere, representations of scene category in object-selective areas were not facilitated by objects. This was further confirmed by a significant interaction (*F*(2,36) = 6.29, *p* = 0.004) between category-selective areas in the left hemisphere (scene-selective, object-selective) and context (scene, scene with object, object).

**Figure 5:**
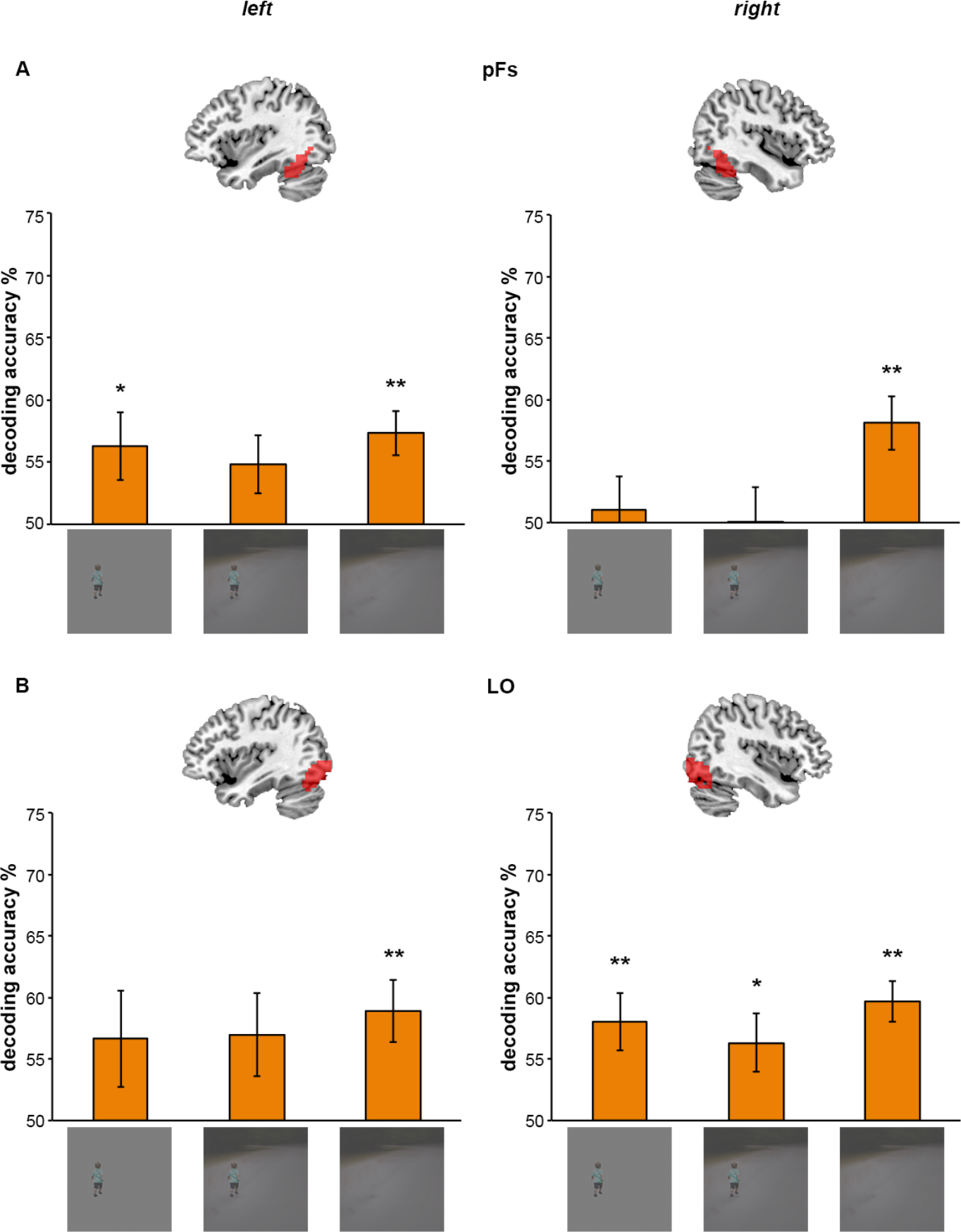
Cross-decoding scene category in object-selective areas. Object-selective areas (red regions in brain maps) were defined by stronger responses to objects than scrambled objects in a separate localizer run: (A) Posterior Fusiform gyrus (pFs); (B) Lateral Occipital cortex (LO). Results show no differences in decoding accuracy for degraded scenes, degraded scenes with object and isolated object in object-selective areas. Thus, objects did not facilitate the classification of degraded scenes in object-selective areas. Data are represented as mean distance from chance (50% decoding accuracy) ±SEM. \**p*<0.05. \*\**p*<0.01.

## Discussion

In the current study, we examined the contribution of contextual object cues to the neural representation of real-world scenes. The key finding was that objects affected scene representation in left, but not right, scene-selective areas, with no difference between effects in OPA and PPA. The magnitude of contextual facilitation in the left scene-selective areas was significantly correlated with scene category information carried by objects presented on gray background of mean luminance of the original scene. In addition, objects did not facilitate scene representations in EVC and object-selective areas. These results provide neural evidence for interactions between object and scene processing, with object processing facilitating scene representations in left scene-selective areas.

The basis for interpreting the current findings is that the dependent measure in all conditions was scene layout classification (indoor/outdoor). Within this framework, we show that objects, which are not an integral feature of scene layout, provide information sufficient to facilitate the representation of scenes in left scene-selective areas, but not right scene-selective areas. This suggests an interaction between object and scene processing, with object processing informing scene representations in left PPA/OPA. This raises the question of whether object information was processed within the scene-selective cortex, or fed to it by external regions. Taking into account the well-established functional dissociation between object- and scene-selective processing (Dilks et al., 2013; Ganaden et al., 2013; Harel et al., 2013; Mullin & Steeves, 2011; Park et al., 2011), we conclude that object information is most likely processed in object-selective cortex and thereafter relayed to scene-selective cortex for contextual facilitation. This is also in line with our previous study, which provided evidence for the reverse interaction, with scene processing in scene-selective areas informing object representation in object-selective areas (Brandman & Peelen, 2017).

What appears to speak against this interpretation is the finding that the response to objects alone allowed for above-chance decoding of the layout of the scene from which the objects were taken (Figure 2). This raises the possibility that both objects and scenes are processed in (left) scene-selective areas. It should be noted, however, that the objects in the isolated-objects condition were still presented in their original scene position, were presented in an experimental context of scenes, and were overlaid on a background frame that had the mean luminance of the corresponding scene. In some of these cases, the object subjectively still evokes the percept of a scene, or at least its coarse layout, with the gray background simply being a more extremely degraded scene (e.g. traffic light and painting in Figure 1a). As such, we believe that the above-chance scene decoding in the isolated-objects condition may reflect the same interactive process as in the scenes-with-objects condition, though to a lesser extent. The strong positive correlation between these two effects is in line with this interpretation.

What is it about the object that facilitates scene category? We propose two plausible explanations. The first, following from the idea that left scene-selective representations are more semantically abstracted, is that the object provides semantic context. By this account, an object more likely to appear in an outdoor scene would bias the representation to open scene layouts, whereas an object likely to appear indoors would imply a closed layout. The second mechanism by which objects may facilitate scene representation is via a fill-in effect. As such, the objects may assist in directly disambiguating the 3D spatial layout by providing relative cues of dimension and distance, thereby facilitating not just semantic interpretation but rather the visual percept of scene layout. Further research is needed to dissociate the contributions of each of these proposed mechanisms.

Regarding the debate about the representational content of scene-selective areas, we hypothesized that areas encoding high-level information about scene layout would exhibit higher discriminability of degraded scenes presented with objects than without, whereas areas that rely exclusively on information provided by scene-typical visual features would not benefit from contextual object cues (Figure 1c). The current findings provide evidence for both high-level views (Aminoff, Kveraga, & Bar, 2013; R. A. Epstein, 2008; Kravitz et al., 2011; Park et al., 2011; Wolbers et al., 2011) and low-level views (Nasr et al., 2014; Nasr & Tootell, 2012; Rajimehr et al., 2011; Zeidman et al., 2012) of scene-selective areas. Particularly, representations of scene category in the left PPA and OPA were facilitated by the presence of an object. This suggests that contextual cues increase the amount of information used to disambiguate open and closed scenes, even when low-level typical scene features are similar. Thus, scene-selective areas in the left hemisphere carry high-level representations of scene layout, beyond information carried by global visual scene features. Such representations may contribute to proposed roles of scene-selective areas in high-level functions such as navigation (R. A. Epstein, 2008) and semantic context processing (Aminoff et al., 2013). In contrast, in the right hemisphere, the PPA and OPA did not appear to represent inferred scene layout, but rather represented scene category based only on global visual scene features present in the image. Together, these findings demonstrate different roles for left and right PPA/OPA in the representation of scene layout.

Why are high-level representations of scene layout left lateralized? One possibility is that representations of scenes in the left hemisphere are more highly abstracted. Given the extensively-studied role of the left hemisphere in semantic processing (Price, 2010), and following the traditional left-verbal/right-visuospatial model, this may suggest that the representations of scenes in the left PPA and OPA are more closely related to their semantic interpretation, which is facilitated by contextual object cues, whereas scene representations in the right PPA and OPA are independent of semantic cues and therefore unaffected by objects. In addition, our results may also be in line with an alternative lateralization model, by which the left hemisphere is specialized for high-frequency information and the right hemisphere is specialized for low-frequency information, due to hemispheric differences in receptive-field sizes (Sergent, 1983). Following this idea, left scene-selective areas may be more sensitive to high-frequency object cues, whereas right scene-selective areas would be better tuned to global low-frequency features. Both these hypotheses should be tested in future studies examining the lateralization of semantic representation and sensitivity to objects in the PPA and OPA.

To conclude, we have found that objects play an important role in the processing of real-world scenes. Specifically, our results show that the representation of scene layout in PPA/OPA was facilitated by contextual object cues. Intriguingly, this effect was strongly left lateralized, demonstrating separate roles for left and right PPA/OPA in the representation of visual scenes, whereby left PPA/OPA represents inferred scene layout, influenced by contextual object cues, and right PPA/OPA represents a scene’s global visual features.

